# A Gradient-based Spectral Explainability Method for EEG Deep Learning Classifiers

**DOI:** 10.1101/2021.07.14.452360

**Authors:** Charles A. Ellis, Mohammad S.E. Sendi, Robyn L. Miller, Vince D. Calhoun

## Abstract

The automated feature extraction capabilities of deep learning classifiers have promoted their broader application to EEG analysis. In contrast to earlier machine learning studies that used extracted features and traditional explainability approaches, explainability for classifiers trained on raw data is particularly challenging. As such, studies have begun to present methods that provide insight into the spectral features learned by deep learning classifiers trained on raw EEG. These approaches have two key shortcomings. (1) They involve perturbation, which can create out-of-distribution samples that cause inaccurate explanations. (2) They are global, not local. Local explainability approaches can be used to examine how demographic and clinical variables affected the patterns learned by the classifier. In our study, we present a novel local spectral explainability approach. We apply it to a convolutional neural network trained for automated sleep stage classification. We apply layer-wise relevance propagation to identify the relative importance of the features in the raw EEG and subsequently examine the frequency domain of the explanations to determine the importance of each canonical frequency band locally and globally. We then perform a statistical analysis to determine whether age and sex affected the patterns learned by the classifier for each frequency band and sleep stage. Results showed that δ, β, and γ were the overall most important frequency bands. In addition, age and sex significantly affected the patterns learned by the classifier for most sleep stages and frequency bands. Our study presents a novel spectral explainability approach that could substantially increase the level of insight into classifiers trained on raw EEG.

## I. Introduction

The frequency domain of electroencephalograms (EEG) is a rich source of information on brain function and has played a vital role in EEG analyses over previous decades. Many studies have used traditional machine learning approaches to gain insight into EEG data. These studies have often used spectral features and standard explainability methods or interpretable classifiers [1]. The growth of deep learning has presented new opportunities for EEG analysis and automated feature extraction [2]. However, the use of automated feature learning with raw EEG data complicates the problem of explainability.

Training neural networks on raw data makes explainability difficult because typical deep learning explainability methods like layer-wise relevance propagation (LRP) [3] find the importance of each time point [4]. An importance value for each time point is too complicated to draw useful conclusions. There is thus a great need for approaches to gain insight into spectral features learned by deep learning models trained on raw EEG.

Strategies to do this include: (1) interpretable classifiers and (2) post-hoc explainability methods. Interpretable classifiers use specialized filters that restrict the domain of learnable features to only spectral features [5][6]. This restriction is problematic if one desires to learn more than just spectral features. Post-hoc approaches do not have the same restrictions as interpretable classifiers [7]–[9]. However, they have key shortcomings. (1) Most approaches have used frequency domain perturbation. (2) Most approaches are global, not local, explainability methods.

Perturbation methods can create out-of-distribution samples that may prevent them from accurately explaining a classifier [10]. All but one approach has used frequency domain perturbation [7]–[9]. In that study, the authors combined activation maximization and LRP [9]. Global methods show the relative importance of each frequency band to the overall performance of the classifier. Local methods explain the importance of each frequency band to the classification of individual samples. They can also be used to form a global estimate of feature importance [11]. Local methods have a key advantage over global methods. They can be used for insight into how demographic and clinical variables affect the patterns learned by the classifier for different frequency bands and different classes [11]. Most post-hoc spectral explainability studies have used global approaches [7], [8]. Only one study has presented a local approach [12].

Gradient-based feature attribution (GBFA) methods offer an alternative that addresses the shortcomings of existing studies [13]. GBFA methods provide local explanations and do not perturb samples. They give importance values for each point in an input sample based upon the importance of the features that those points compose. As such, periodic features like sinusoids of a particular frequency should have reoccurring explanations, and an analysis of the frequency domain of the explanations should indicate the relative importance of each frequency.

In our study, we present a novel spectral explainability approach that addresses the key shortcomings of existing methods. We train a convolutional neural network (CNN) for automated sleep stage classification. We apply LRP, a popular GBFA method and perform a Fast Fourier Transform (FFT) of the explanations to estimate frequency band importance. We then perform a statistical analysis of the local spectral explanations to determine whether participant age and sex affected the patterns learned by the classifier for each sleep stage and canonical frequency band.

## II. Methods

Here we describe the data, preprocessing, classifier, explainability approach, and statistical analyses of our study.

### A. Description of Data

We used the Sleep Cassette subset of the Sleep-EDF dataset [14] on Physionet [15]. The dataset was originally used in [16] to examine the effects of age and sex upon sleep. It had 153 approximately 20-hour recordings from 78 participants (41 female and 37 male). Participants varied in age from 25 to 101 years with a mean (μ) of 58.79 years and a standard deviation (σ) of 22.15 years. We analyzed the EEG Fpz-Cz electrode. The data were recorded at a sampling frequency of 100 Hertz (Hz). Technicians annotated each 30-second segment as *Unmarked, Movement, Awake*, Rapid Eye Movement (*REM*), non-REM 1 (*NREM1*), *NREM2, NREM3*, and *NREM4*.

### B. Description of Data Preprocessing

We separated each recording into 30-second samples. We removed *Unmarked* and *Movement* samples and removed all *Awake* samples from the start of each recording. We combined *NREM3* and *NREM4* into *NREM3* [17]. To ensure that *Awake* had the same number of samples as *NREM2* (i.e., the other majority class), we removed some *Awake* samples at the end of the recordings. We z-scored each recording. Our final dataset had 215,668 samples and was imbalanced with *Awake, NREM1, NREM2, NREM3*, and *REM* composing 39.43%, 9.98%, 32.05%, 6.05%, and 22.98% of the data, respectively.

### C. Description of Classifier

We used a CNN architecture developed in [18]. Figure 1 shows the architecture. We used 10-fold cross-validation with training, validation, and test sets having 63,7, and 8 randomly assigned participants, respectively. We used categorical cross- entropy loss with class-based weighting to account for data imbalance. We used Adam with an adaptive learning rate starting at 0.001 and decreasing by an order of magnitude after each five epochs without an increase in validation accuracy. To assess classifier performance, we computed the μ and σ of the precision, recall, and F1 score for each class across folds.

**Fig. 1.**
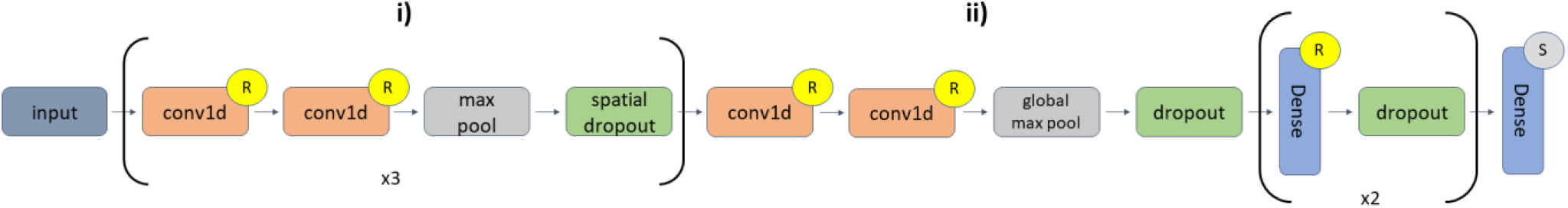
CNN Architecture. **i)** had 6 1D convolutional (conv1d) layers. The first repeat had two conv1d layers with 16 filters and a kernel size of 5, a max pooling layer with a pool size of 2, and a spatial dropout layer with a rate = 0.01. The second repeat had two conv1d layers with 32 filters and a kernel size of 3, a max pooling layer with a pool size of 2, and a spatial dropout layer with a rate of 0.01. The third repeat had 2 conv1d layers with 32 filters and a kernel size of 3, max pooling with a pool size of 2, and spatial dropout with a rate of 0.01. In **ii**), the last two conv1d layers had 256 filters with a kernel size of 3 and were followed by global max pooling and dropout with a rate of 0.01. The first dense layer had 64 nodes and a dropout layer with a rate of 0.1, and the second dense layer had 64 nodes and a dropout layer with a rate of 0.05. The last dense layer had 5 nodes. Layers with an “R” or “S” were followed by ReLU or Softmax activation functions, respectively.

### D. Description of Explainability Approach

We used LRP in our study [3]. LRP was first developed for image analysis but has been applied to electrophysiology [4]. We used the Innvestigate toolbox to implement LRP [19]. In LRP, a sample is fed to a classifier and the output node for the top class is assigned a total relevance value of 1. Based upon relevance rules, portions of that relevance are iteratively assigned to each previous layer until they are propagated to the input layer of the network. Relevance can be positive or negative. Positive and negative relevance identify features that support the classification of a sample as its assigned class and as a class other than its assigned class, respectively. We used 2 relevance rules: the ε-rule and αβ-rule. The ε-rule has a parameter (ε) that filters smaller relevance values during propagation when increased. The αβ-rule has parameters α and β that control the portion of positive and negative relevance that are propagated, respectively. We used the ε-rule with values of 0.01 and 100 and the αβ-rule (α = 1, β = 0). After applying LRP to each raw EEG test sample, we performed an FFT of the relevance assigned to each sample and averaged the power of the relevance assigned to each frequency bin. We used δ (0 – 4 Hz), θ (4 – 8 Hz), α (8 – 12 Hz), β (12 – 25 Hz), and γ (25 – 50 Hz). The values for each particular frequency band reflected the relative importance of each frequency bin. We displayed the local results for each sample over time for the ε-rule (ε = 100). We also computed the mean absolute relevance for each frequency band and classification group across folds to approximate the global importance of each frequency band.

### E. Description of Statistical Analyses

To examine the effects of sex and age upon patterns learned by the classifier for each sleep stage and correct classification group, we trained an ordinary least squares regression model with age and sex as independent variables and the relevance of a frequency band as the dependent variable. This enabled us to control for the effects of each variable when examining the relationship of the other variable with the local explanations. We analyzed the explanations for the ε-rule (ε = 100). We repeated the regression analysis for each frequency band. We then used false discovery rate correction with the sex p-values and with the age p-values to account for multiple comparisons (α = 0.05).

## III. Results and Discussion

We describe and discuss the results for the spectral LRP and statistical analyses. We also discuss the study limitations and potential future work.

### A. Performance Results

Table 1 shows the CNN’s performance results. The classifier performed most effectively for *Awake* and least effectively for *NREM1*. This is interesting given that *Awake* and *NREM1* are the largest and nearly the smallest classes, respectively. *NREM1* performance may have also been particularly low because *NREM1* and *REM* are similar and have been combined into a single class in some previous studies [20]. The model performed well for *NREM2, NREM3*, and *REM*, obtaining best F1 and precision for *NREM2* and best recall for *NREM3*.

**TABLE I.** Classification Performance Results

|  | Awake | NREM1 | NREM2 | NREM3 | REM |
| --- | --- | --- | --- | --- | --- |
| **F1** | 91.28±02.53 | 41.94±03.07 | 77.38±03.32 | 66.59±11.32 | 68.71±05.97 |
| **Precision** | 96.82±01.44 | 36.10±05.23 | 84.97±04.09 | 57.45±11.33 | 64.88±10.75 |
| **Recall** | 86.45±04.27 | 52.04±08.26 | 71.66±07.23 | 80.66±14.96 | 74.57±02.58 |

### B. Global Approximation of Frequency Band Importance

Figure 2 shows the results for our global approximation of frequency band importance. Overall, δ, β, and γ seemed to be the most important frequency bands for the correct classification groups. For correctly classified *Awake*, the results differed slightly across LRP rules. For the ε-rule (ε = 0.01 and 100), γ was the most important frequency band, though γ was not as important as δ for the αβ-rule. Both the ε-rule (ε = 100) and αβ- rule identified δ as more important than β, and all three relevance rules identified γ as more important than β. For *NREM1*, all three relevance rules identified γ as most important, and the ε-rule (ε = 0.01 and 100) identified β as more important than δ. The ε- rule (ε = 100) and αβ-rule identified δ as the most important frequency bin for *NREM2*, and all three relevance rules found β more important than γ. For *NREM3*, the ε-rule (ε = 100) and αβ- rule found δ to be most important. The ε-rule (ε = 0.01) and αβ- rule found β to be second most important for *NREM3*. Interestingly, the ε-rule (ε = 100) indicated that θ might have been more important than γ. For *REM*, γ was the most important frequency band for the ε-rule (ε = 0.01 and 100). Based upon the ε-rule (ε = 100) and αβ-rule, δ was the second most important frequency band for *REM*. β was the third most important, followed by θ and α. It is probable that instances in which the ε- rules disagreed with the αβ-rule that there were particularly large amounts of negative relevance, and in instances where the ε-rule (ε = 100) and αβ-rule disagreed with the ε-rule (ε = 0.01), it is likely that the noisiness of ε-rule (ε = 0.01) was problematic.

**Fig. 2.**
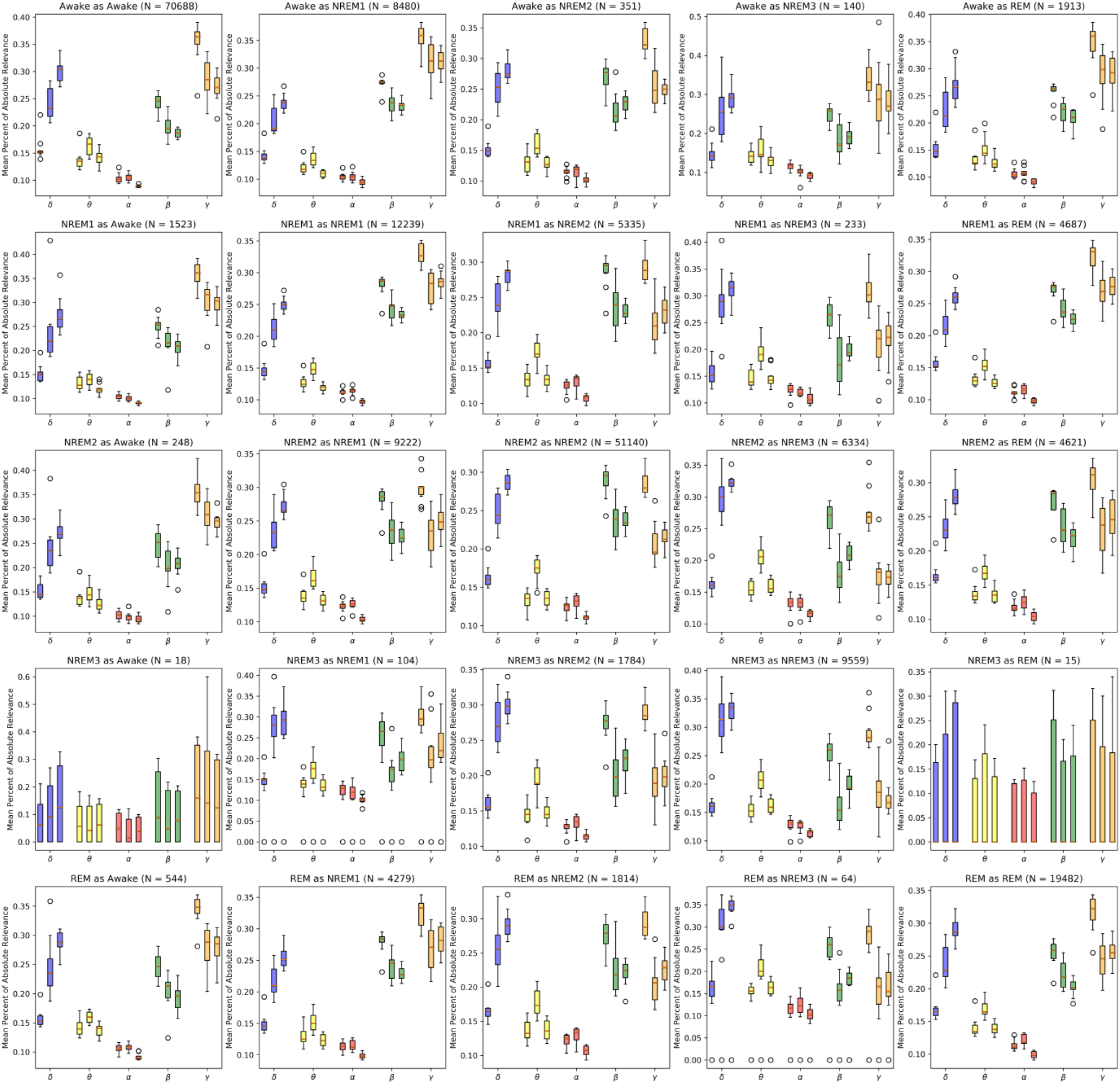
Global Approximation of Frequency Band Importance. Plot shows results for all folds. From left to right in each group of boxes are results for the LRP ε-rule (0.01), ε-rule (100), and α-β-rule. The title of each panel shows the number of samples in each classification group. Correct and incorrect classification groups are shown in the panels on the left-to-right diagonal and off the left-to-right-diagonal, respectively. The y-axis is the mean absolute power of the relevance. The x-axis indicates the frequency bands analyzed. Interestingly, δ, β, and γ were the most important frequency bands overall.

Across classes incorrect classification groups had slight differences relative to correct classification groups in how the mean absolute power of the relevance was distributed across folds for nearly all frequency bands. In some cases, incorrect classification groups had different levels of variance in their importance distributions across folds (e.g., the ε-rule with ε = 0.01 and 100 for *NREM3* classified as *NREM2* in γ and δ in *REM* classified as *Awake*). In instances, where incorrect classification groups had higher variance than correct classification groups, that could indicate that the incorrect patterns learned related to a particular class varied across folds while the correct patterns remained more consistent across folds.

### C. Frequency Band Importance Over Time

Figure 3 shows the importance of each frequency band for a 120-minute segment of a recording from Participant 7. The importance of some frequency bands seems to vary throughout the recording. The δ importance increased from 0 to 60 minutes, held a consistent level of peak importance from 60 to 80 minutes during *REM*, decreased again at around 85 minutes, increased up to around 110 minutes, and decreased again. Most of the increases in δ importance were associated with increases in δ activity. In contrast, γ began with a peak level of importance that waned from 0 to 20 minutes during the transition from *NREM1* to *NREM2*. Early in the recording θ transitioned from a state of low importance to a state of oscillating relatively high and low importance. The α importance was consistently low. The β importance started at higher levels and remained high until *NREM3* onset at around 45 minutes.

**Fig. 3.**
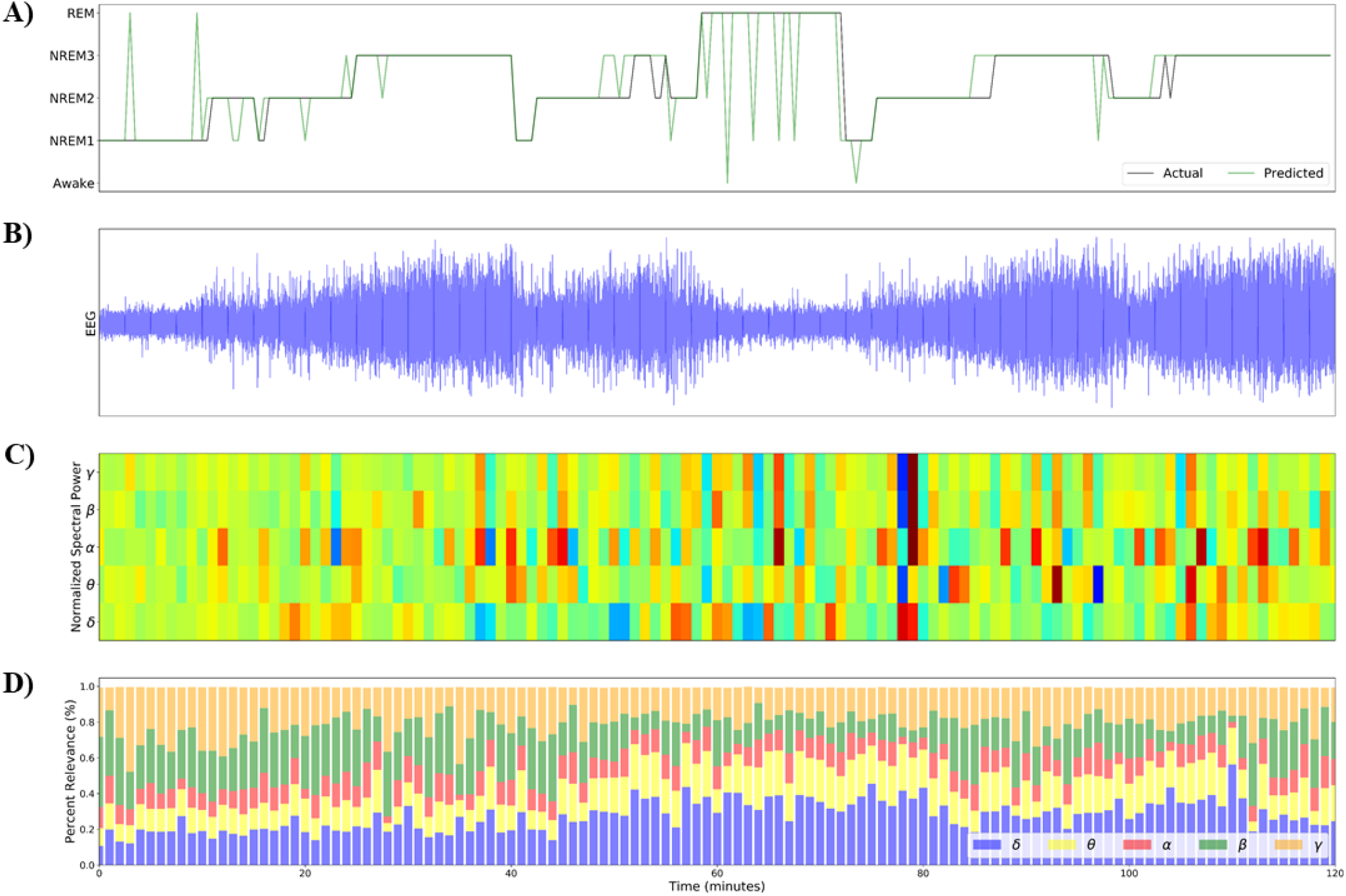
Frequency Band Importance Varies over Time. This figure shows the spectral explainability results over a 2-hour period from Subject 7. Panel A shows the actual and predicted classes of each 30s sample throughout the 2-hour window. Panel B shows the raw EEG. Panel C shows the mean spectral power for each frequency band of each 30-second sample. Panel D shows the importance for each frequency band over time in a normalized manner (i.e., divided by maximum spectral value) that enable an easier visualization of the variation of the spectral power over time. Panel D shows the portion of the power of the relevance assigned to each frequency band. Panels A and D have legends that assist with figure interpretation.

### D. Effects of Age and Sex On Local Explanations

Figure 4 shows the results for the statistical analysis that we performed to identify the effects of age and sex on the local explanations. Most frequency bands and classification groups had significant relationships with age and sex, which is not unexpected given that the dataset was collected to study the effects of age and sex upon sleep [16].

**Fig. 4.**
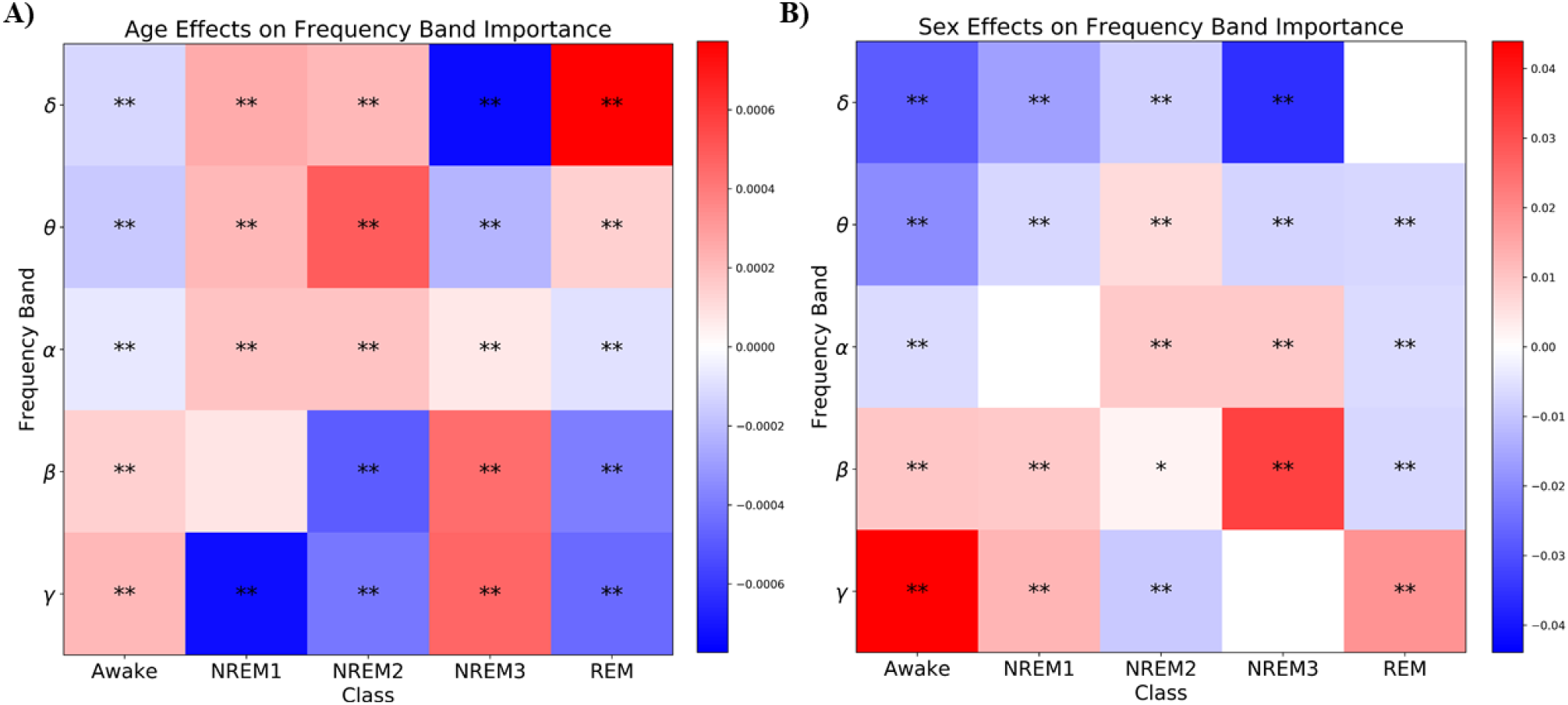
The Effect of Age and Sex on Frequency Band Importance for ε = 100. Panels A and B show the effects of age and sex on frequency band importance, respectively. The x-axis of each panel shows the class corresponding to each column, and the y-axes show the frequency band corresponding to each row. The heatmaps indicate the coefficients resulting from the regression analysis. White blocks indicate frequency bands and classes that are not significantly affected by the demographic variable (i.e., p > 0.05). Colored blocks indicate coefficients with corresponding significant p-values (p < 0.05) following correction. Asterisks * and ** indicate coefficients with corresponding significant p-values at p < 0.01 and p < 0.001 after FDR correction, respectively. For age, a red coefficient indicates that importance for that frequency band and class increased with age. For subject sex, a red coefficient indicates that importance for that frequency band and class was higher in males than females.

For age, a red coefficient value in Figure 4 indicated that importance for that frequency band and class increased with age. Age had a mix of positive and negative relationships with frequency band class importance. The largest magnitude effects of age were found for δ, β, and γ, though age did have a moderate effect on θ for *NREM2*. Interestingly, in *NREM1*, γ importance decreased with age, which corresponded with more importance for all other frequency bands with increased age. For *NREM2*, β and γ importance decreased with age, which corresponded with increased importance for all other frequency with age. *NREM3* δ importance also decreased with age, while β and γ importance increased with age. In contrast, REM δ importance increased with age while β and γ importance decreased with age.

For sex, a red coefficient in Figure 4 indicated that males had more importance for that frequency band and class than females. Sex had a mix of positive and negative effects upon frequency band importance across classes. The strongest effects were in δ, β, and γ and θ to a lesser degree. There were some instances in which sex seemed to have no significant effect (e.g., *REM* δ, *NREM1* α, and *NREM3* γ). In a few cases, frequency bands demonstrated opposite changes in importance based upon sex.

Males had more *Awake* γ importance than females and less δ and θ than females. Males also had more *NREM3* β than females but less δ than females. In some minor cases, there were corresponding changes in frequency band importance (e.g., *NREM1* δ and θ importance was greater in females while β and γ importance were greater in males).

### E. Limitations and Future Work

In this study, we applied three LRP relevance rules individually. Using different relevance rules for different parts of the network might improve the explanation quality [21]. Additionally, other GBFA methods that might provide better explanations [22]. For our statistical analysis of the effects of age and sex upon the classifier, we only examined correct sample groups. Future studies might examine the effects of age and sex upon explanations for incorrect classification. While the focus of our paper was on our novel explainability approach rather than our classifier, other deep learning architectures have obtained higher classification performance than the architecture that we used. The use of state-of-the-art classifiers might enable the use of our approach for automated biomarker identification.

## IV. Conclusion

In this study, we trained a classifier for automated sleep stage classification and presented a novel approach for local spectral insight into deep learning classifiers trained on raw EEG data. In contrast to the majority of existing approaches that require the ablation or modification of the spectral domain of data and risk creating out-of-distribution samples, our approach avoids that problem altogether. We then performed a statistical analysis that enabled us to examine the effects of age and sex upon the patterns learned by the classifier for each frequency band and sleep stage. Our results indicated that δ, β, and γ were the most important frequency bands across most sleep stages. We further found that age and sex significantly affected the patterns learned by the classifier for most sleep stages and frequency bands. Our study offers a novel approach with the potential to significantly improve the insight that can be gained for classifiers focused on EEG classification.

